# Coordinated Head Direction Representations in Mouse Anterodorsal Thalamic Nucleus and Retrosplenial Cortex

**DOI:** 10.1101/2022.08.20.504604

**Authors:** Marie-Sophie H. van der Goes, Jakob Voigts, Jonathan P. Newman, Enrique H. S. Toloza, Norma J. Brown, Pranav Murugan, Mark T. Harnett

## Abstract

The sense of direction is critical for survival in changing environments and relies on flexibly integrating self-motion signals with external sensory cues. While the anatomical substrates involved in head direction (HD) coding are well known, the mechanisms by which visual information updates HD representations remain poorly understood. Retrosplenial cortex (RSC) plays a key role in forming coherent representations of space in mammals and it encodes a variety of navigational variables, including HD. Here, we use simultaneous two-area tetrode recording to show that RSC HD representation is nearly synchronous with that of the anterodorsal nucleus of thalamus (ADn), the obligatory thalamic relay of HD to cortex, during rotation of a prominent visual cue. Moreover, coordination of HD representations in the two regions is maintained during darkness. We further show that anatomical and functional connectivity are consistent with a strong feedforward drive of HD information from ADn to RSC, with surprisingly little reciprocal drive in the corticothalamic direction. Together, our results provide direct evidence for a concerted global HD reference update across cortex and thalamus, and establish the underlying functional connectivity that supports this coordination.

## Introduction

In order to enable efficient navigation, internal representations of self-location and orientation must be updated as sensory experience and behavioral demands fluctuate. Changes in environmental information are known to trigger remapping of place (Muller, and Kubie, 1987), grid tiling (Fyhn, et al., 2007) and HD (Taube, Muller and Ranck, 1990; Goodridge *et al*., 1998; Knierim, Kudrimoti and McNaughton, 1998). In the insect and mammalian HD systems, remapping has been observed in the form of rotations of preferred firing directions (PFD) of HD cells in response to rotations of a prominent visual cue - often a color-contrast card (Taube, Muller and Ranck, 1990; Goodridge *et al*., 1998), a narrow band (Seelig and Jayaraman, 2015) or scene (Kim *et al*., 2019) on an LED screen. Hebbian synaptic plasticity mechanisms, acting in specific circuit arrangements, have been proposed to explain these phenomena in the fly ellipsoid body (Fisher, et al., 2019; Kim, et al., 2019). Network models with similar architecture have been applied to the rodent HD system (Hahnloser, 2003; Knight, et al., 2014a; Page, et al., 2014; Skaggs, et al., 1995). However, unlike the fly brain (Franconville, et al., 2018; Hanesch, et al., 1989), the circuitry that drives HD remapping in the rodent brain is not yet resolved. Several studies have indicated that cortical regions play a role in the stability and the visual anchoring of HD (Clark, et al., 2010; Golob, and Taube, 1999; Goodridge, and Taube, 1997), but whether the HD representation is first aligned to the sensory cues in cortex and then updated in downstream regions is still unknown. Understanding the dynamics and the connectivity of visual-HD integration is the first necessary first step to uncover the mechanisms that lead to remapping in the mammalian HD system.

Previous studies in the rat indicate that the HD code in the Anterodorsal Thalamic Nucleus (ADn), the necessary thalamic relay of HD to the hippocampal formation and cortex (Calton, et al., 2003; Frost, et al., 2021; Goodridge, et al., 1997; Jenkins, et al., 2004; Winter, et al., 2015), becomes unstable and is less likely to remap to reflect cue rotations after lesions to the Retrosplenial Cortex (RSC) (Clark, et al., 2010) or the Post-Subiculum (POS) (Goodridge, et al., 1997). Both of these regions are strongly interconnected with visual areas (Sugar, et al., 2011; Van Groen, and Wyss, 2003), with each other (Kononenko, and Witter, 2012; Wyss, and Van Groen, 1992), and with ADn (Jankowski, et al., 2013). While HD dominates the spatial code in POS (Taube, Muller and Ranck, 1990; Peyrache, Schieferstein and Buzsáki, 2017; Laurens *et al*., 2019), RSC exhibits diverse visuo-spatial activity (Cho, and Sharp, 2001; Fischer, et al., 2019; Mao, et al., 2017; Powell, et al., 2020; Voigts, and Harnett, 2020a), with complex receptive fields shaped by multiple spatial correlates (Alexander, et al., 2020; Alexander, and Nitz, 2017, 2015; Jacob, et al., 2017). As a cortical association area, RSC plays a critical role in spatial cognition (Knight, and Hayman, 2014b; Mitchell, et al., 2018; Vann, et al., 2009) and spatial memory (Miller, et al., 2019, 2014): rats and humans with RSC lesions show impairments in route planning as well as identification and flexible use of navigational landmarks (Hindley, et al., 2014; Maguire, 2001; Pothuizen, et al., 2008; Vann, and Aggleton, 2004). Specifically, in the intact RSC, associations between egocentric and allocentric reference frames become evident with spatial tasks (Alexander, et al., 2020, 2015; Shine, and Wolbers, 2021; van Wijngaarden, et al., 2020). These kinds of computations might underlie the emergence of navigational landmarks (Auger, et al., 2012; Fischer, et al., 2019; Jacob, et al., 2017; Page, and Jeffery, 2018), as sensory stimuli that appear in the egocentric view and, having been deemed reliable, are ultimately mapped to an abstract representation of space (Barry, and Burgess, 2014; Bicanski, and Burgess, 2016; Yan, et al., 2021). Altogether, this evidence strongly implicates RSC in the integration of visual orienting cues and HD. However, it is unknown how the ensuing changes in HD representation are coordinated between cortical and sub-cortical circuits, and what the role of RSC is in this process. We sought to answer this question by performing simultaneous single unit recording in RSC and ADn in freely moving mice while a visual cue was either rotated around the arena or turned off.

## Results

### Differential encoding of HD in RSC and ADn

To monitor single unit activity in RSC and ADn we implanted independently movable tetrodes assembled in a lightweight microdrive (Voigts, et al., 2020b) targeting the two regions simultaneously in 9 mice. We additionally recorded from RSC alone in two mice, and used carbon fiber electrodes to record from ADn in one more mouse (see Supplementary Fig. 1A for electrolytic lesions in ADn and RSC for each mouse). During the recordings, mice could freely roam in a dark circular arena of 50 cm diameter, inside a large sealed box. The only visual cue was provided by an illuminated subset of LEDs spanning an angle of 20° which formed a wider circle outside and above the arena. 48% of the units we recorded under these conditions in ADn met our criterion for HD cells, whereas 20% in RSC did so (Fig. 1B,C). Our HD cell selection method relied on the amount of directional information and magnitude of the resultant from the tuning curve or the von Mises fit (LaChance, et al., 2022) to the largest peak for multi-peak units against those obtained from shuffling the spikes (Supplementary Fig. 2A). RSC displayed a modest HD code, in agreement with previous findings in rats and humans (Chen, et al., 1994a; Cho, et al., 2001; Shine, et al., 2016, 2021). Directional information in RSC was generally lower than in ADn (bits/spike, median: ADn HD=0.0567, n=225; RSC HD=0.0134, n=204; ADn NonHD=0.0142, n=248; RSC NonHD=0.0074, n=805; Kruskal-Wallis test p <0.0001, p<0.001 for multiple comparisons, except for ADn NonHD and RSC HD cells, where p=0.0675) (Supplementary Fig. 2B). Consistent with previous findings (Taube, 1995), both ADn and RSC HD cells were positively modulated by angular velocity (AV) (cue-on median ADn=1.20, n=225; RSC=1.26, n=204; cue-off median ADn=1.15, n=150; RSC=1.21, n=57; Supplementary Fig. 2G), calculated as the ratio of the firing rate for high (>30°/s) over smaller (<30°/s) angular velocity. However, both during cue-on and -off conditions RSC HD unit firing rates were more sensitive to angular velocity than ADn (Mann-Whitney test p<0.001 and p<0.01 for cue-on and cue-off, respectively). The differences between these two regions were consistent with the higher degree of multi-modal selectivity in RSC (Alexander, et al., 2020, 2015; Cho, et al., 2001; Laurens, et al., 2019), where multiple spatial correlates, contexts and states are mixed with and influence HD coding.

**Figure 1:**
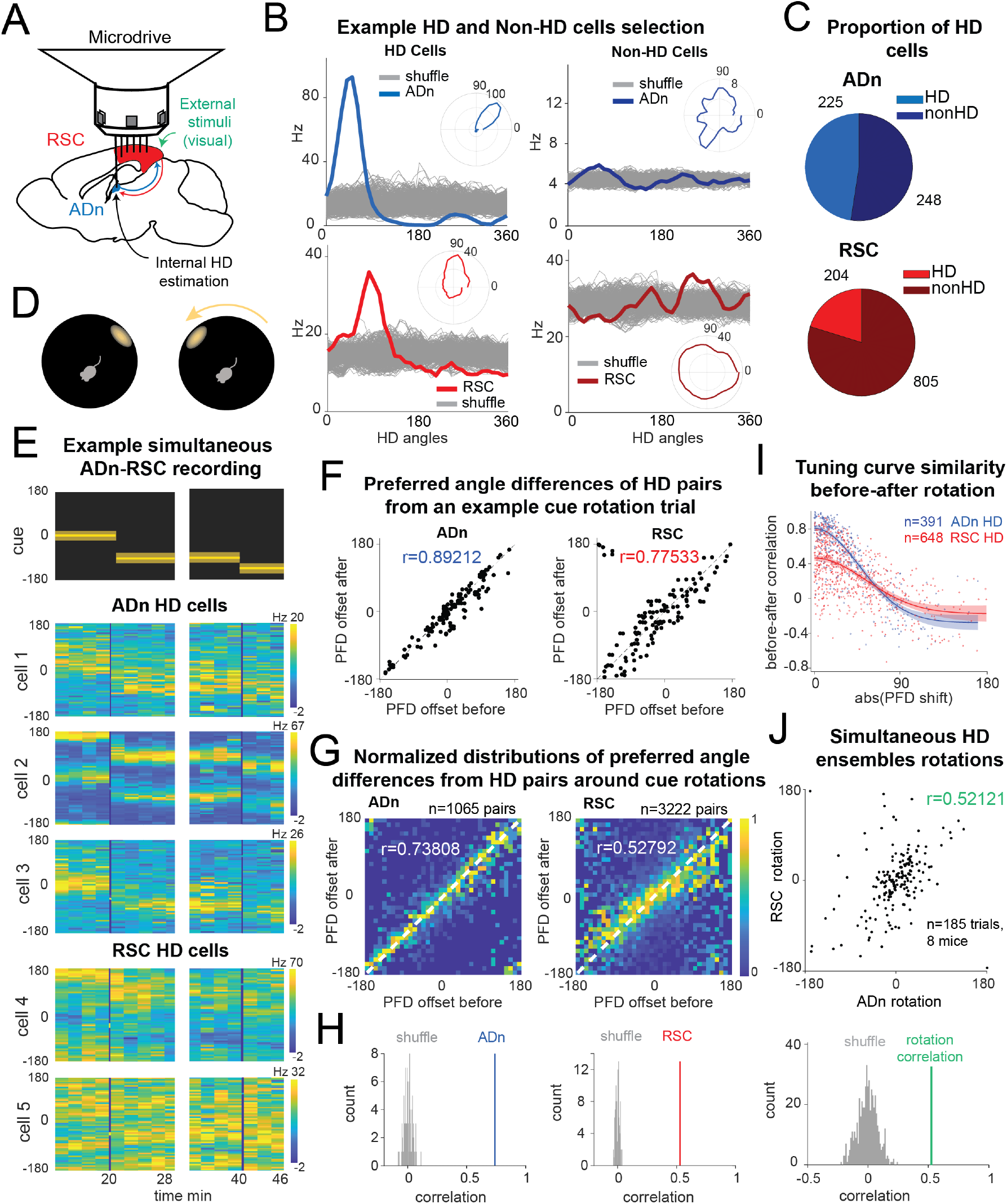
Congruent HD response to visual cue rotation in ADn and RSC, despite differences in strength of HD coding. **A:** Schematic of simultaneous ADn (blue) and RSC (red) tetrode recording. **B:** Tuning curves of examples of HD (left) and non-HD (right) cells in ADn and RSC. Grey lines are the tuning curves obtained from 500 shuffles of the cells firing rates. Insets show the tuning curves in polar coordinates. **C:** Pie charts showing that 48% of cells in mouse ADn meet the HD selection criterion, but only 20% in RSC do (right) (N=12 mice, 8 with simultaneous ADn and RSC, 2 ADn only and 2 RSC only). **D:** Schematic of the arena with the only prominent LED cue before (left) and after 90-degree rotation (right). **E:** Simultaneous ADn-RSC recording from a session where the cue (top) was rotated first by 90° (first segment) and then 45° (second segment). Tuning curves (2 min bins) over time of HD cells in the two regions shift the preferred firing direction (yellow bins, maximal firing rate) in response to the cue rotation. **F:** Scatter plots of preferred firing directions (PFD) differences from all ADn HD cells pairs (left) and RSC HD cells pairs (right) from example trials before versus after rotation. **G:** 2D histograms of PFD differences before versus after rotation from all distinct HD unit pairs from rotation trials of ADn recordings (left) and RSC recordings (right) (ADn correlation: 0.738, N=90 trials, 1065 pairs, 10 mice; RSC correlation: 0.527, N= 91 trials, 3222 pairs, 10 mice). Each column bin is normalized by its maximum value to equally represent all PFD offsets (see Supplementary Fig. 2F for the unnormalized graph). **H:** Correlation values of ADn pairs (left) and RSC (right) from the data in G are above the 99^th^ of 100 randomly drawn angle differences. **I:** Correlation values between before and after rotations HD tuning curves in ADn and RSC plotted against their absolute rotation value. While ADn correlation values sharply decrease with larger PFD rotations, this relationship is less marked and exhibits higher variability in RSC. Lines and shaded areas represent the best gaussian fit to the data and the 95% CI, respectively (R^2^= 0.8, n=321 for ADn and R^2^=0.4, n=648 for RSC). Data points in I are individual neurons from the same rotation trials as in G. **J:** Top, mean rotations from HD cells in ADn vs mean rotations of simultaneously recorded HD cells in RSC. Bottom, correlation value from the data in the left (N=185 trials, 8 mice) is above the 99^th^ percentile of 500 times shuffled rotation trial indices for each HD cell.

### Congruent HD responses to visual cue rotations in ADn and RSC

To challenge the mice’s sense of orientation and determine whether ADn and RSC similarly update the HD frame in response to changes to visual stimuli, we instantaneously rotated the LED cue around the arena by either 45° or 90° (Fig. 1D), in trials ranging from 5 to 40 minutes. Previous work with cue card rotations indicates that HD cells in ADn rotated their preferred directions coherently (Yoganarasimha, et al., 2006), but it remains unclear if the same applies to RSC. Unlike POS, which presents mostly homogeneous, cue-guided rotations of HD fields (Taube, et al., 1990b), diverse responses of RSC HD cells to cue manipulations have been reported (Chen, et al., 1994b), suggesting differential influence of idiothetic and allothetic cues. A recent study in medial entorhinal cortex reported firing rate changes and subsets of HD cells with persistent preferred tuning and others with cue-following tuning to cue manipulations (Kornienko, et al., 2018). In contrast, despite the complexity of an environment with two distinct spatial reference frames, entorhinal HD stays internally organized and unitary, but not consistently anchored to a global reference (Park, et al., 2019). To further investigate whether RSC HD ensembles stay coherent in our behavioral setting, we calculated the angle offsets between the tuning curves of all unique simultaneous HD pairs before and after cue rotation. As expected, the preferred direction difference between pairs of ADn HD neurons remained rigid after cue rotations (Fig. 1F,G,H, circular correlation=0.738, n= 90 trials, 1065 pairs, 10 mice). Despite the reduced HD information and lower resultant in HD cells in RSC compared to ADn (Supplementary Fig 2C&E, median resultant length ADn HD cells: 0.159, non-HD cells: 0.055, RSC HD cells 0.076, non-HD cells 0.041, p<0.0001 for multiple comparisons after Kruskal Wallis test, p<0.0001), we observed a smaller but significant correlation (0.527, n=91 trials, 3222 pairs,10 mice) in preferred direction difference of RSC HD pairs between before and after cue rotations (Fig. 1F,G,H). Similar results were obtained when all rotation trials were included (correlation ADn= 0.7811 n=211 trials, 4686 pairs; RSC 0.432, n=222 trials, 17579 pairs).

The reduced rigidity of RSC HD ensembles could emerge from changes in tuning, namely the degree to which they encode HD, and/or from ensemble variability in preferred direction changes. We quantified the change in tuning, also indicated as “HD score”, as the ratio between the resultant length of the tuning curve before and after the rotation. We found that while both groups had HD scores centered around 1 (ADn median=0.99, n=391, RSC median=0.94, n=648), RSC HD units had slightly more unstable tuning (Kolmogorov-Smirnov p<0.0001, Supplementary Fig. 2H, top). Moreover, the correlation between ADn HD tuning curves before and after rotation was high for small shifts in preferred direction and, as expected, this relationship sharply dropped for larger rotations (Fig. 1I). RSC exhibited more variability, but nonetheless displayed a similar drop in the tuning correlation with large shifts (gaussian fit n=391 R^2^=0.8 for ADn, n=648 R^2^=0.4 for RSC, Fig. 1I). Altogether, we conclude that RSC HD units, despite the higher variability in the tuning, also exhibit generally coherent HD remapping.

To compare how both regions responded to cue rotations, we calculated the mean preferred direction shifts for RSC and ADn HD ensembles. Despite a bias toward the inertial (self-motion-driven) HD estimate, evident through the higher density of rotations around 0, the rotations from the simultaneously recorded ADn and RSC HD ensembles were more correlated (coefficient=0.52, n=185 trials, 8 mice) than those produced by random shifting of RSC and ADn neural activity around the cue rotations (Fig. 1J). This suggested that the HD map is well-locked between the two regions in our behavioral paradigm, despite the differences in HD encoding between ADn and RSC.

### Synchronous shifting of HD representations in ADn and RSC in response to cue rotations

Based on the tuning curves of our HD cells, we concluded that RSC and ADn encode the same HD reference, independent of whether they shift with cue rotation or disregard the cue as an orienting landmark (Fig 1J). However, it was not clear if ADn and RSC also coordinate at a finer temporal scale in response to the cue rotation. To answer this question, we applied a decoding approach to infer the HD representation from the entire recorded population at a high temporal resolution (20 ms bins) in ADn and RSC. We implemented a linear-Gaussian generalized model (GLM) that related ADn or RSC ensemble neural activity to HD obtained from behavioral tracking (Supplementary Fig. 3A). For each trial we estimated the weight coefficients based on the stable cue period before rotation, excluding the last 80 seconds, and tested the model on the remaining stable period, by calculating the difference between the decoded HD and the HD from headstage tracking. On average 75% of the decoded errors from the test period were below 60.133° ± 3.44 in ADn (n=303) and 72.46° ± 3.44 in RSC (n=311) (p<0.0001, Mann-Whitney test) (Supplementary Fig. 3B,C). However, given that HD is modulated by angular velocity (Supplementary Fig. 2G, top), we considered whether the decoding accuracy was affected by these state changes. Indeed, decoding performance dropped for high compared to low angular velocity (Supplementary Fig. 3D&E), (low angular velocity means: ADn=56.8°, RSC=68.46°, high angular velocity means: ADn=71.67°, RSC=90.83°). Two-way ANOVA revealed significant differences between the two regions and also between the two velocity states, but only a small effect for the interaction between the two variables (p=0.045), suggesting that the angular velocity effect was largely similar between the two regions.

The errors from our GLM decoder reflected the mean rotations of the ensemble HD neurons tuning curves both from ADn and RSC (Supplementary Fig. 3F&G, 0.546 and 0.485 circular correlation coefficients for ADn, n=277, and RSC, n=271, respectively, p<0.0001 for both) suggesting that our method largely captured the changes in neural activity with cue rotation. The mean decoded errors after cue rotations from simultaneously recorded ADn and RSC ensembles were also correlated (Fig. 2B, 0.664 circular correlation coefficient, n=213), confirming the findings from the mean tuning curves rotations (Fig. 1J). Similarly, we also observed a high density of decoded rotations values around zero. This effect likely resulted from devaluation or bias toward internal HD estimates after multiple exposures to visual and internal HD mismatches within the arena (Knierim, et al., 1998).

To investigate whether the egocentric experience of the cue influenced the rate of under-rotations, we compared rotations occurring when the cue was well outside of the visual field of the mouse before rotation (Dräger, and Olsen, 1980; Sterratt, et al., 2013) or not (>154° or >-154°, calculated at the center of the cue, with 0° aligned to the mouse’ snout; Supplementary Fig. 4A). We found that, despite an overall effect of the size of the rotation (p<0.001 ANOVA with 3 factors, p>0.05 for interactions with other factors or effect of region grouping), there were no detected differences between the groups (multiple comparison after Bonferroni correction p>0.05, cue out of view: small rotation n=26 trials, ADn 0.31, RSC 0.26, big rotation n=18 trials, ADn -0.127, RSC 0.0569; cue in view: small rotation n=69 trials, ADn 0.35, RSC 0.32, big rotation n=100 trials, ADn -0.001, RSC 0.194, Supplementary Fig. 4B). This suggested that even though large mismatches were more likely to result in under-rotations, we could pool the results of the rotation analyses. Furthermore, we did not find any correlation between the size of the decoded rotation in ADn and RSC and the egocentric bearing of the cue before (Supplementary Fig. 4C, 0.043 and 0.006 correlation coefficients respectively for ADn, n=303, and RSC, n=311) or after rotation (Supplementary Fig. 4D, 0.015 and -0.090 correlation coefficients respectively for ADn and RSC). Altogether, these analyses indicate that, in our behavioral setting, the initial egocentric experience of the cue rotation was not a factor in the under-rotations of the HD representation.

Next, we asked how changes in environmental stimuli alter the rate of HD reference shifts, and if this differ across brain regions. By applying our decoding strategy, we first observed that decoded errors drifted to cue-set offsets at variable speed (Fig. 2C). On average we observed similar, and mostly slow, HD reference shifts in both ADn and RSC (Supplementary Fig. 5B). Interestingly, we observed closely matched trajectories to the new HD offsets in simultaneously recorded neurons from both ADn and RSC (Fig. 2C). To quantify if initial jitter in the drift of the HD representations were indicative of a region-specific response to the new angular position of the cue, we calculated the temporal cross-correlations between simultaneous ADn and RSC decoded errors immediately after cue rotations (75 s window). To isolate the effect of successful update of the HD reference, we included only trials where the absolute mean HD shift was at least 11.5° or above. While substantial trial-to-trial variability in the correlation level was observed, the majority of trials had peaks at 0 ms time lags (two-sample Kolmogorov-Smirnov test between data and null distribution, p<0.0001 for both stable and shifted), both before and after cue rotations (Fig. 2D,E) (Wilcoxon Signed-Rank test p=0.4117, n=159 trials). Similar results were obtained for shorter decoded error traces immediately after cue rotation (25 s window), albeit with higher variability (Supplementary Fig. 5A, two-sample Kolmogorov-Smirnov p<0.0001 for both stable and shifted; Wilcoxon Signed-Rank test between shifted and stable p=0.38, n=159 trials). This confirmed that the two regions were coordinated even in the initial stages of the shifting of the HD representation. Contrary to our predictions (i.e. that RSC would lead the HD reference update via visual integration of the cue’s new angular position), we found that the two regions were highly synchronized in this regard. Variability was observed in the histograms from the cross-correlations after cue rotation, but they lacked any specific bias for anticipatory or delayed time lags. Changes in the neural activity, and therefore in HD decoding, could emerge following the cue rotation as a response to visual stimuli, potentially explaining the slightly decreased synchrony compared to stable cue periods (Fig. 2E).

**Figure 2:**
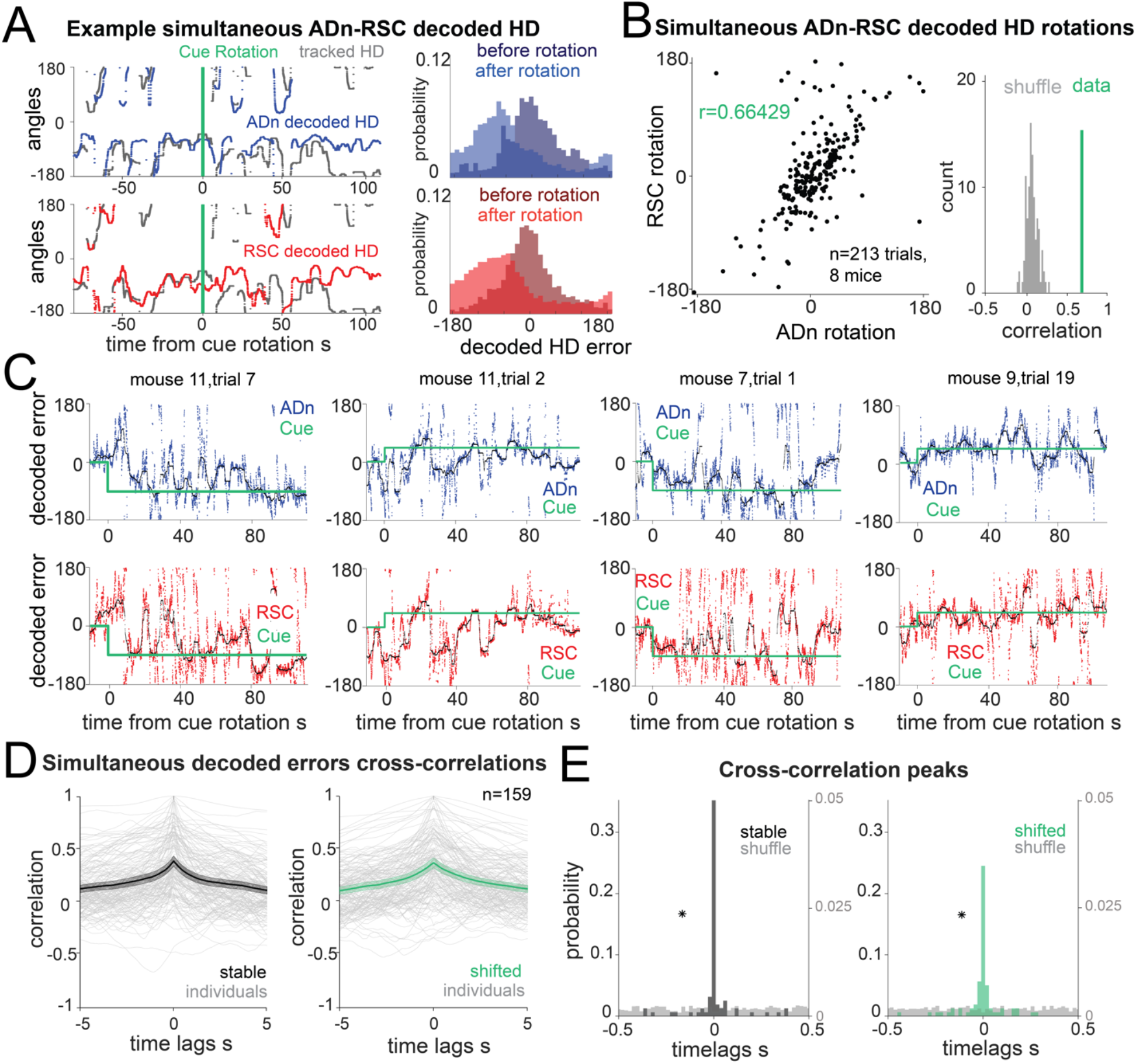
Synchronous shifting of ADn and RSC HD representation in response to cue rotation. **A**: Left, Example of decoded HD in ADn (top, blue line) and RSC (bottom, red line) from a simultaneous recording in the two regions before and after cue rotation (yellow line at t= 0 s). Grey line, mouse HD from tracking. Right, probability-histograms of the difference between the tracked and the decoded HD shown on the left in ADn (top, blue) and RSC (bottom, red). Darker shades, decoded error before rotation, lighter shades after rotation. **B:** Left, decoded ADn vs paired RSC rotation (N=213 trials, 8 mice). Right, correlation coefficient between the decoded ADn and RSC rotation (green) is above the 99^th^ percentile of the 100 times randomly shifted RSC decoded HD for each trial (grey). **C:** Four examples of paired ADn (top row) and RSC (bottom row) decoded HD errors drifting toward the target (yellow). Black lines, median-smoothed error over a 10 s window. The first example is the decoded error of the traces shown in A. **D:** Mean and 95% confidence intervals of temporal cross correlation between paired decoded errors before rotation (left, black) and immediately after rotation (right, green) (75 s long segments, N=159 out of 204 paired trials with mean ADn rotation >11.5°, 8 mice). Grey, individual trials. **E:** View in the -0.5 s to 0.5 s range of the probability normalized histograms (20 ms bins) of the time lags corresponding to the peak correlation values from all trials in E, left, before rotation, right, after rotation. Left y-axis scaled to show the uniformity of the null distributions (grey). Asterisks indicate the real distributions are significantly different from null (two-sample Kolmogorov-Smirnov test, p<0.0001 for both stable and shifted). No difference between stable and shifted trial correlations was observed (Wilcoxon Signed-Rank test p=0.4117).

### Correlated HD drift in darkness in ADn and RSC

Visual cues influence the HD signal in ADn and POS by providing an external anchoring reference (Taube, et al., 1990b; Taube, and Burton, 1995), counteracting the drift from stochastic error in angular velocity integration observed in darkness (Mizumori, and Williams, 1993; Stackman, and Taube, 1997; Valerio, and Taube, 2012). RSC is also necessary for path integration-based navigation in darkness (Elduayen, and Save, 2014), likely by integrating motor (Yamawaki, et al., 2016) and angular velocity signals (Hennestad, et al., 2021; Keshavarzi, et al., 2021) together with the incoming HD from thalamic nuclei and POS, to form a representation of orientation. However, it is unknown if visual cues are necessary for maintaining the coordination of HD representations in ADn and RSC. To resolve this issue, we challenged the sense of orientation in a subset of mice by turning the LED off after a period of cue-on baseline (Fig. 3A). HD cells maintained the same initial preferred directions while the cue was on in a stable position, but were more variable while the cue was off, and sometimes continuously drifted during prolonged darkness (Fig. 3B). On average, we observed modest levels of drift in darkness (Supplementary Fig. 6A, p<0.05 Kuiper test between total cue-on and cue-off mean drift distribution, n=72 trials in ADn). When we applied the same decoding strategy (Fig. 3C and Supplementary Fig. 6D) to quantify the drift in the two regions, we observed correlated HD representations between ADn and RSC both during cue-on and cue-off periods. This is evidenced by the strong diagonal band across the column-normalized 2D histograms of the HD decoded errors (Fig. 3D left). A small but significant effect on the HD score change between cue-on and cue-off in RSC compared to ADn (Supplementary Fig. 2H bottom, ADn median=0.96, n=150, RSC median=0.82, n=71, p=0.014 Mann-Whitney test) suggested that darkness does affect the tuning of RSC HD neurons to some extent (Chen, et al., 1994b). However, the circular correlations of the decoded drifts were similar between cue-on (r=0.33318) and cue-off periods (r=0.31128) and were significantly higher than the correlations with shuffled RSC drifts (Fig. 3D, right).

**Figure 3:**
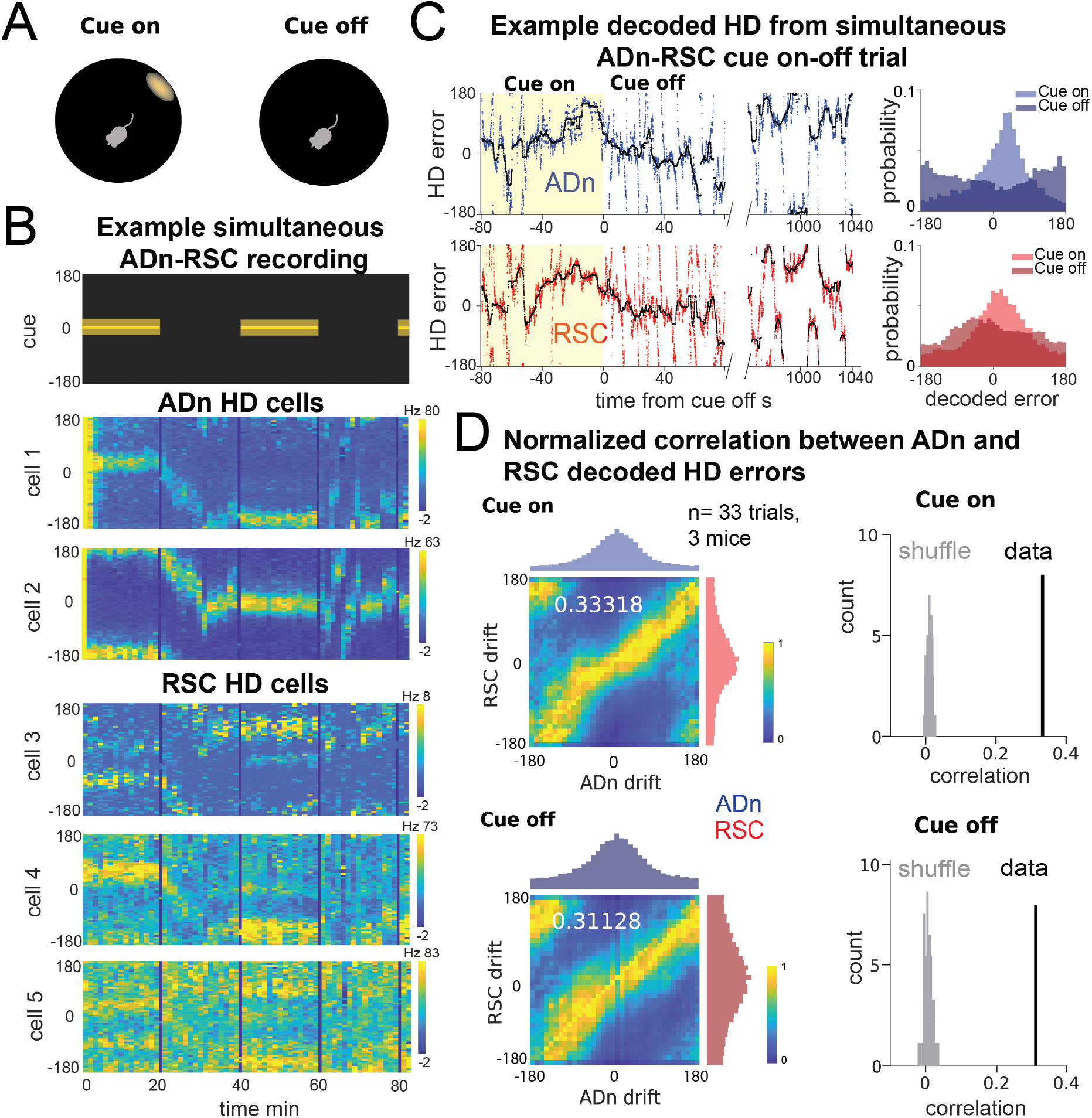
Correlated HD drift in darkness in ADn and RSC. **A:** Schematic of cue-on/off trials. **B:** Simultaneous ADn-RSC recording from a session where the cue (top) was turned on and off. **C:** Left, simultaneous ADn (blue) and RSC (red) decoded HD errors from the first cue-on (yellow shaded) and -off trial of the example shown in B. Black, median-smoothed decoded error over a 5 s window. Right, probability normalized histograms of the decoded HD error from the example in C in ADn (blue, top) and RSC (red, bottom). The lighter shaded histograms are from the cue-on segments, the darker shaded from the cue-off. **D:** Top, left: 2D histogram of simultaneous ADn and RSC decoded errors from cue-on, normalized by the maximum column value per bin (i.e. in the ADn dimension); above and on the side, marginal distributions of ADn and RSC drift, respectively. Top, right: correlation value (green) from the not-normalized data in the left is above the 99^th^ percentile of the distribution obtained after randomly shifting the RSC drift in each trial (grey), both for the cue-on (top) and cue-off (bottom) segments. Bottom, left: same 2-D histogram and marginal distributions during cue-off; right: correlation of the real data vs shuffle. (N=33 trials, 3 mice).

In the awake behaving rodent, angular velocity drives the update of the ongoing HD, which, in the absence of prominent visual cues, can drift inconsistently from the current HD reference (Skaggs, et al., 1995; Stackman, et al., 2002; Valerio, and Taube, 2016). Moreover, RSC HD neuron firing rates are more sensitive to angular velocity modulation both during cue-on and during cue-off periods thank ADn (Supplementary Fig. 2G, p<0.01 Mann-Whitney test). How this variable affects the unanchored HD in ADn and RSC is not known. We therefore also examined the effect of angular velocity on the simultaneous ADn-RSC HD drift in darkness. We found that even under the challenges of greater angular velocity (>30°/s), more often associated with larger HD errors during cue-on (Supplementary Fig. 3D&E), the coordination between the two regions is maintained (Supplementary Fig. 6B&C, high angular velocity cue-on r=0.29 versus cue-off 0.28, low angular velocity cue-on r=0.39 vs cue-off r=0.36). Our results therefore indicate that the HD representation is also coordinated between ADn and RSC in the absence of a visual input and during periods of HD instability.

### Asymmetry in the RSC-to-ADn and ADn-to-RSC connectivity

Temporal coordination of HD representation on the order of 20 ms or less across different structures could be accomplished by direct monosynaptic connections or by concurrent input from different areas, particularly for the visual update of the HD reference. To investigate the anatomical substrate for direct connectivity between ADn and RSC, we performed retrograde monosynaptic rabies tracing (Wickersham, et al., 2007) experiments in ADn and RSC. We found that RSC cells from the same subregion where we performed tetrode recordings (Supplementary Fig. 1A,B) received dense anterior thalamic inputs, and particularly ADn (Fig. 4A). Conversely, ADn exhibited surprisingly sparse presynaptic RSC cell labeling. These cells were frequently localized in the granular and ventral portion of RSC (Fig. 4B). These results are consistent with previous studies in the rat, which reported reciprocal but similarly asymmetric connectivity between ADn and RSC (Shibata, 1993, 1998; Van Groen, and Wyss, 1990).

**Figure 4:**
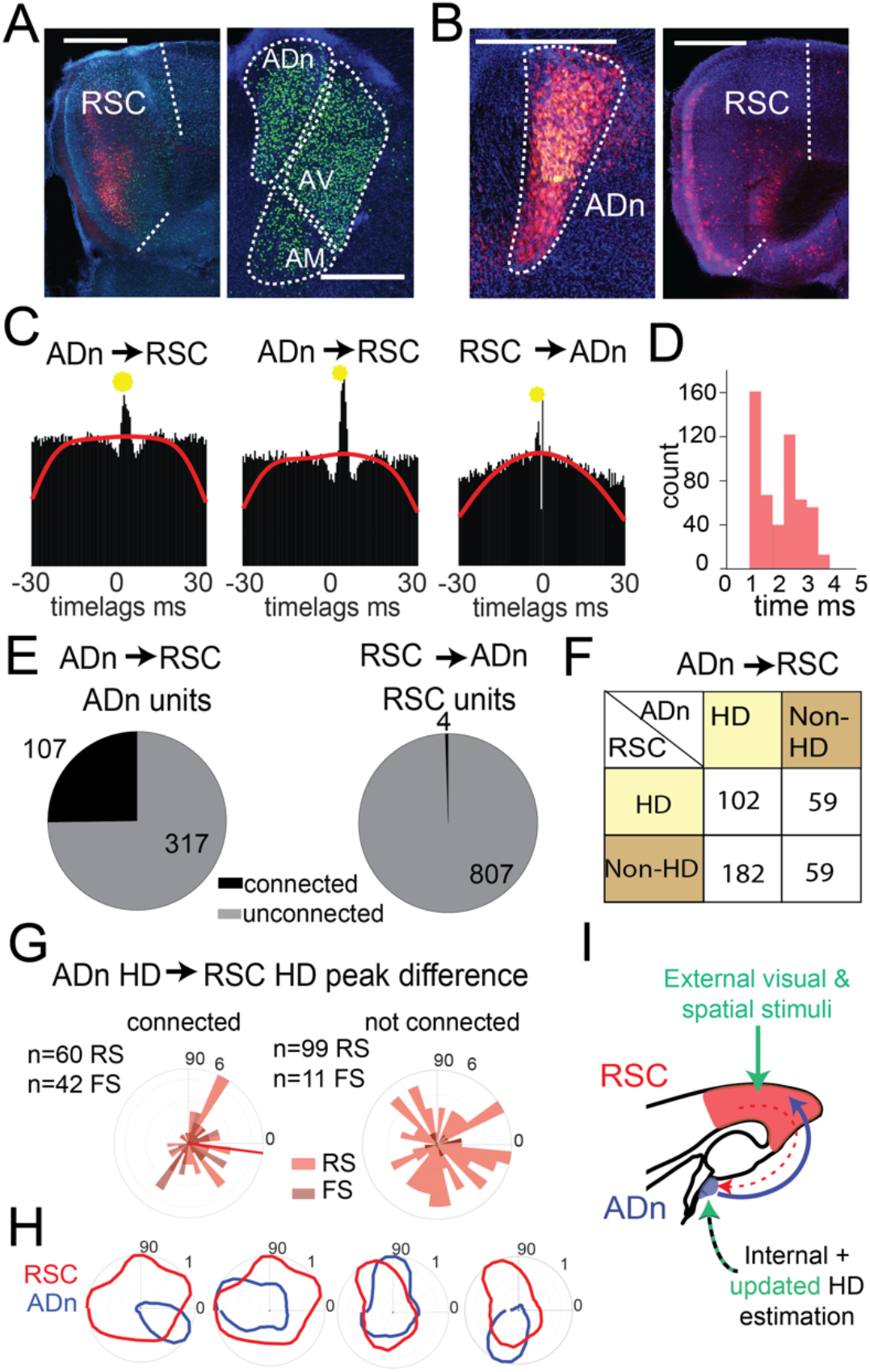
Asymmetric connectivity between RSC and ADn. **A:** Monosynaptic rabies tracing of inputs to RSC (left, starter cells in red) shows a high density of presynaptic cells in ADn (right, green). **B:** Monosynaptic rabies tracing of inputs to ADn (left, starter cells identified by the overlap of blue, green and red) shows a low density of presynaptic cells in RSC (right, red) and mostly in A29. Scale bar in A and B, 0.5 mm. **C:** Examples of cross-correlograms with putative excitatory connections (yellow circle) from ADn to RSC (first 2 examples) and RSC to ADn (third example), showing a sharp peak between 1 and 5 ms time lag above the baseline (red line) at more than 99.9% of the cumulative Poisson distribution. **D:** Distribution of the latencies of the peaks in the cross-correlograms for ADn-to-RSC connections. **E:** Number of ADn units with putative connections to RSC (left) and of RSC units with functional connections to ADn (right). **F:** Breakdown into HD-and non-HD coding of the putative pre- and post-synaptic partners of connected ADn-to-RSC pairs. **G:** Polar plot distributions of the differences between preferred directions of connected ADn HD units and their putative HD-coding synaptic partners in RSC (N=102 units, 60 RS and 42 FS, 6 mice, left) and the same ADn HD units and all other HD-coding non-synaptic partners (N=110, 99 RS and 11 FS, 6 mice, right). Red line on the left plot indicates the circular mean (−7.44°) of the RS peak differences; Rayleigh test for non-uniformity p=0.0086 for the RS synaptic partner, p<0.001 for RS and FS, p=0.1290 of the non-synaptic partners (right plot). **H:** Polar plots of tuning curves of 4 example pairs of connected ADn to RSC HD units with variable preferred directions. **I:** Adapted schematic from Fig. 2.3 showing that the connectivity from RSC to ADn is nearly absent and that the visually-guided updates in the HD frame emerge from a strong feedforward HD input from ADn.

We next examined whether the HD coordination we observed in our recordings could emerge from ensembles in RSC or ADn that were functionally connected and could convey updated or visually-anchoring HD information. To this end, we performed spike cross correlation between all possible ADn-RSC pairs, considering the spikes occurring during cue-on periods. Putative monosynaptic connections were identified in the cross correlograms as sharp peaks above the baseline (Fig. 4C) between 1 and 5 ms from the time of ADn unit firing (Fig. 4D). Using this metric (Peyrache, et al., 2015; Stark, and Abeles, 2009), we identified 6.4% of all possible pairs with a monosynaptic connection in the ADn-to-RSC direction (522 out of 8083) and only 0.08% in the RSC-to-ADn direction (7 out of 8083). In terms of unit counts, 107 out of 424 ADn units had at least one connection to RSC, while only 4 out of 811 RSC units had a connection with ADn (Fig. 4E). The ADn-to-RSC connectivity was divergent, with a mean number of RSC synaptic partners of 4.64 for ADn HD cells and 3.33 for non-HD cells. When we focused on the connectivity during darkness, we found similar results: 45 out of 226 units in ADn (3 mice) had at a connection, while no RSC units were connected. The generally lower connectivity rate was likely due to the reduced sample size of cue-off trials, but suggested that even in the absence of visual cues corticothalamic projections were not a substrate for HD coordination. ADn’s dense connectivity largely originated from identified HD cells (284 connected pairs from ADn HD cells versus 118 from non-HD ADn cells) and included both HD as well as non-HD partner cells in RSC (Fig. 4F). HD-coding was a feature of both RS and FS units in RSC (Supplementary Fig. 7), and both groups received ADn connections (Fig. 4G, left plot).

Finally, the distribution of the preferred direction differences between ADn HD-coding units and their HD-coding synaptic partners in RSC showed a small but significant bias toward similar tuning (Fig. 4G left polar plot, n=102 units, 60 RS and 42 FS, 6 mice, Rayleigh test for non-uniformity p=0.0086 for the RS synaptic partners, circular mean - 7.44°). This similarity was not observed for the preferred direction differences between the same ADn units and all other non-synaptically connected HD-coding units, whose distribution was uniform (Fig. 4G right polar plot, n=110, 99 RS and 11 FS units, p=0.1290 Rayleigh test for the RS units). Together, these results suggest that ADn sustains the RSC HD code with a widespread feedforward connectivity to both RS and FS units, a connectivity that targets not only clearly HD-tuned units, directly shaping their preferred directions (Fig. 4H), but also units with more complex, presumably multimodal, receptive fields. On the other hand, the very sparse RSC-to-ADn connections alone are unlikely to drive the change or the stability in the presence of visual cues, in HD preferred directions (Fig. 4I).

## Discussion

Our data show that the HD representation in ADn and RSC is closely coordinated, both in conditions when visual cues are stable and during adaptation to a new reference in response to cue rotation (Fig. 2D,E). This is also true for HD drift in darkness (Fig. 3D), showing that visual input is not necessary for maintaining these coordinated representations, and that other sensory modalities, such as angular velocity, optic flow, and/or motor efference copy, influence ADn and RSC HD. Finally, functional connectivity data further indicate that RSC is likely not wired to drive visual reference updates, or even maintain visual anchoring, via direct corticothalamic input to ADn. ADn, however, provides strong feedforward input to RSC, likely driving the local HD code there (Fig. 4E-H). Together, our results provide direct evidence against the hypothesis that visually-guided updates in the HD reference would first appear in RSC, as a result of visual integration, and then be conveyed to ADn. We conclude that the visually-driven updating of HD is a more complex process that privileges coordination across brain regions over sustained error signals with mismatched representations.

Using a simple generalized linear encoding model of HD (Supplementary Fig. 3A) we decoded HD at a fine temporal scale (Supplementary Fig. 3B,C) with ensembles of 4 to 30 units ADn and 8 to 90 in RSC. We observed variable and mostly slow drifts of the HD representation to a new target (Fig. 2C and Supplementary Fig. 5B). These drifts suggest that an experience- and time-dependent weighting of the internal HD estimation against the egocentric bearing of the cue takes place as the new HD reference is determined. Repeated exposure to an unstable cue is known to cause landmark devaluation (Knierim, Kudrimoti, Mcnaughton, 1995; Knight *et al*., 2014) and, together with extended navigation within the same environment, increased the incidence of under- or 0° rotations in our data (Supplementary Fig. 4B). In addition, these conditions may affect the speed of these shifts in our paradigm. Slow, continuous drifts of the HD representation after cue rotation as reported here (Fig. 2C, Supplementary Fig. 5B) have also been observed in flies (Kim, et al., 2017) and in rat (Knierim, et al., 1998) and mouse (Ajabi, et al., 2021) ADn, with the exception of one study in the rat ADn (Zugaro, et al., 2003) where immediate HD shifting was observed in specific cue-heading configurations. While behavioral and arena-configuration differences may underlie this discrepancy, it remains to be resolved if the cause of such slow drifts lies in: 1) the configuration of the polarizing visual stimulus, including the distance between the cue and the mouse, with farther away cues being less impacted by egocentric view and having more control over HD (Zugaro, et al., 2001); 2) the memory of previous experiences of cue rotations (Ajabi, et al., 2021; Knierim, et al., 1995); and/or 3) the intrinsic time course of synaptic plasticity associated with the learning of the new landmark orientation (Goodridge, et al., 1998; Kim, et al., 2019, 2017; Page, et al., 2014; Skaggs, et al., 1995; Yan, et al., 2021).

Our anatomical and functional connectivity experiments reveal a striking asymmetry between the strong feedforward ADn-to-RSC HD drive (Fig. 4E-G) and the sparseness of RSC-to-ADn connections. This asymmetry was more extreme than that observed in previously reported ADn-POS connectivity (Peyrache, et al., 2015; van Groen, and Wyss, 1990), but similar to anatomical tracing in the rat (Shibata, 1998), and possibly further exacerbated by the widespread sampling of RSC locations in our recordings (i.e. granular vs dysgranular, L2/3 vs L5; Supplementary Fig.1). Furthermore, the general bias toward similar tuning between connected ADn and RSC HD units suggests that ADn HD code might not be simply inherited in RSC, as has been shown in POS (Peyrache *et al*., 2015; Peyrache, Schieferstein and Buzsáki, 2017), but likely integrated with other spatial codes via different functional circuit organizational principles, that include recruitment of FS and RS neurons (Simonnet, et al., 2017).

How would the visual cue integration that anchors HD be reflected in the ensemble representation? We hypothesized that an “error” signal would appear as a temporal offset in the HD of the two regions: specifically, the RSC HD update by integration of visual inputs, would precede that of other regions, in our case ADn. Contrary to this hypothesis, our decoding showed no consistent temporal offset in HD representation during shifting (Fig. 2D,E). Importantly, this was true regardless of the cue rotation and the devaluation of the anchoring effect of the cue, evident from the under- or no rotations (Fig. 1J, 2B and Supplementary Fig. 4). At the same time, even in the absence of visual cues, RSC and ADn were closely coordinated during small and large HD drifts, a phenomenon that has recently been proposed to be mediated by intact cerebellar inputs (Fallahnezhad, et al., 2021). This coordination, which we directly show under two visual challenges, is likely sustained by a strong and widespread feedforward ADn-to-RSC connectivity, where the updated HD reference may already be computed upstream of ADn (Yoder, et al., 2015). This framework is consistent with an existing hypothesis that visual anchoring may compete with and, depending on the manipulation, dynamically bias the memory of the internal estimation from angular velocity (Ajabi, et al., 2021; Knierim, et al., 1998).

Whether distinct circuit mechanisms are recruited to coordinate the learning of the new orienting cue according to the behavioral demands and the complexity of the navigation task is not known. POS, through its reciprocal connections with visual areas, could provide the visual reference information to ADn, RSC and LMN, the obligatory HD path upstream of ADn (Yoder, et al., 2015; Yoder, and Taube, 2011). Another possible route includes the cortico-thalamic control through thalamic reticular nucleus, which readily and densely inhibits ADn and receives presubicular and retrosplenial connections (Vantomme, et al., 2020).

The HD coordination and striking sparseness of RSC-to-ADn connectivity do not preclude, however, that RSC could support the change in HD reference through activation of dedicated ensembles encoding the orienting “landmark” (Bicanski, et al., 2016; Mitchell, et al., 2018; Page, et al., 2018) or with conjunctive HD-visual fields. In fact, RSC is necessary for ADn HD alignment to visual cues (Clark, et al., 2010) and the dense interconnection between RSC and several regions of the hippocampal formation (Sugar, et al., 2011; Wyss, et al., 1992) may support coordinated HD representation across the brain as a mechanism to ensure consistent flexible spatial computations relevant to behavior output. Future experiments using multi-site high-density recordings with laminar probes in RSC could directly assess the activity patterns, at the single unit and population level, associated with the learning of the new cue orientation. Under tight temporal control between cue rotation and the animal view and limited cue devaluation with repeated trials, these experiments could test for the presence of orienting landmark-coding cells.

## Methods

### Behavior and Subjects

All animal procedures were performed in accordance with NIH and Massachusetts Institute of Technology Committee on Animal care guidelines. We used adult (>8 weeks old) C57BL/6 from Charles River and from Jackson Laboratory RRID: IMSR_JAX:000664 and one Vgat-Ires-Cre C57 BL/6 mice (RRID:IMSR_RBRC10723). 4 females and 8 males were used for tetrode recordings, and 2 12-week old mice were used for rabies tracing experiments. Mice were kept on a 12-hour light/dark cycle with unrestricted access to water. 8 of the implanted mice underwent mild (up to a 10% reduction in body weight) food restriction. Of the implanted mice, 8 were housed isolated in conventional cages, 4 with siblings in rat cages with running wheels. One mouse had channelrhodopsin expression in cortical interneurons, but this aspect was not investigated in the present study.

The behavioral arena was 50 cm in diameter with a 25 cm cylinder wall, surrounded by an outer cylinder of 80 cm diameter and 30 cm height, where a string of 132 white LEDs (Adafruit, APA102) covering the upper circumference provided the only light source. The arena was enclosed in a 78×86×84 cm wooden dark box to shield from lighting and noise. In food deprived mice, pellets (Bioserve) were sprinkled on the floor to allow continuous exploration during long recordings. To provide novelty in the environment and induce exploration, two types of arena walls (black pvc with a white paper at the upper edge and opaque clear plastic) were used and changed when the cue rotation did not produce shifts in the HD tuning.

The visual cue was a set of computer-controlled (Teensy 3.2) LEDs spanning an angle of 20° with brightness following a gaussian with peak at the center and sd of 1. 2 weeks after surgery mice were habituated to a single cue or no cue at all while units were monitored. Different starting cue angular positions for the recording sessions were sampled and different sequences of rotations of ±90° and ±45° were played. For recordings with the Open-Ephys ONIX system (Newman, et al., 2019) with the commutator, rotations and cue on-off switches occurred every 20 to 40 minutes, versus the 5 to 20 minutes for recordings with the first generation Open-Ephys system (Siegle, et al., 2017) without commutator. Sessions length varied based on the animals’ behavior, with a minimum of 2 up to 11 rotations/on-off switches. Cue rotations occurred in consecutive “jumps” from one angular position to the next. In 3 mice, periods of darkness were interleaved in some rotation sessions.

### Electrodes and Drive Implants Surgeries

Light weight drives for tetrode recordings were fabricated following the guidelines in (Voigts, et al., 2020b) for a total of 16 independently-movable tetrodes per drive. Arrays were designed to simultaneously target ipsilateral ADn and RSC, for a total length of 2.8 mm and a width of 0.5 mm. To increase the yield of units especially for ADn, some guide tube positions were occupied by two tetrodes. Tetrodes were constructed from 12.7 μm nichrome wired (Sandvik – Kanthal, QH PAC polyimide coated) with an automated tetrode twisting machine (Newman, 2020) and were gold plated to lower the impedance to a final value between 150-300 Ohm. One mouse was implanted with 32 carbon fiber electrodes (∼100 Ohm, (Guitchounts, et al., 2013) in ADn only, whose position was fixed since surgery.

All surgeries were performed using aseptic techniques. Mice were anesthetized with isoflurane (2% induction, 0.75%–1.25% maintenance in 1 l/min oxygen) and secured in a stereotaxic apparatus. Body temperature was maintained with a feedback-controlled heating pad (DC Temperature Control System, FHC). Slow-release buprenorphine (1 mg/kg) and dexamethasone (4mg/kg) were pre-operatively injected subcutaneously. After shaving of the scalp, application of hair-removal cream and disinfection with iodine and ethanol, an incision was made to expose the skull. For implants, after cleaning with ethanol, the skull was scored and a base of dental cement (C&B Metabond and Ivoclar Vivadent Tetric EvoFlow) was applied. A burr hole was drilled over prefrontal cortex close to the olfactory bulb for placement of the ground screw (stainless steel) connected to a silver wire. Sometimes an additional burr hole and ground screw, connected to the other with silver epoxy, provided extra stability. For drive implants with tetrode arrays, a large craniotomy from ∼0.3 to ∼ 3 mm from Bregma, and from the midline to ∼0.95 mm ML at the level of M2 and ∼0.7 mm ML at the level of RSC was drilled. After durotomy, the drive was lowered onto the surface of the brain with one RSC (AP ∼2.400, ML ∼0.150 mm, DV ∼0.200 mm) and one ADn (AP ∼0.350 mm, ML ∼0.975 mm, DV ∼1.800 mm) -targeting tetrodes extended for guiding the placement of the array. For the carbon fibers implant, a smaller (∼1 mm diameter) craniotomy, followed by durotomy, allowed lowering of the bundle of fibers into ADn (AP: 0.68 mm, ML 0.75 mm, DV 2.65 mm). The drive, or the fiber frame, was then secured to the skull with dental cement, the skin incision was partially closed with sutures and the mouse was placed in a clean cage with wet food and a heating pad and monitored until fully recovered. All drive implants were done on the right hemisphere.

### Viral Surgeries

The same stereotactic procedures were applied to viral surgeries. For ADn rabies tracing experiments, a burr hole was drilled over AP 0.68 mm, ML 0.75 mm coordinates and 25 nL of 1:1:1 mixture of helper viruses, pAAV-syn-FLEX-splitTVA-EGFP-tTA and pAAV-TREtight-mTagBFP2-B19G (Wickersham) and AAV2/1.hSyn.Cre (Janelia Farms) was delivered at a rate of 60nL/min through a glass pipette lowered to DV 2.65 mm. This injection was followed by 50 nL of (EnvA)SAD-ΔG-mCherry (Wickersham) two weeks later at the same location, and after 7 days the brains were processed for histology. For RSC rabies tracing experiments, the coordinates were AP 2.8 mm, ML 0.45 mm, DV 0.75 and 0.45 mm, and the injection of 50 nL of 1:1 mixture of AAV2/1.hSyn.Cre(Janelia Farms) and AAV1-hsyn-DIO-TVA66T-dTom-CVS-N2C(g) (Allen Institute) was followed 3 weeks later by a 100 nL of EnvA dG CVS-N2C Histone-eGFP (Allen Institute) before histological processing 9 days later. 5 minutes after each injection, the pipette was slowly withdrawn and the incision was sutured.

### Immunohistochemistry and confocal imaging

Brain fixation with 4% paraformaldehyde in PBS was achieved with transcardial perfusion for monosynaptic rabies tracing experiments and with drop fix for electrolytic lesions retrieval from the drive-implanted mice. After being left overnight at 4°C, brains were sectioned coronally at 100 μm thickness with a floating section vibratome (Leica VT1000s), washed in PBS and then labeled with 1:1000 DAPI solution (62248; Thermo Fisher Scientific). All sections were mounted and coverslipped with clear-mount with tris buffer (17985-12; Electron Microscopy Sciences). Confocal images were captured using a Leica TCS SP8 microscope with a 10X objective (NA 0.40) and a Zeiss LSM 710 with a 10x objective (NA 0.45). ML and DV coordinates for cortical tetrode rotations (Fig. 1) were measured in ImageJ/FIJI (National Institutes of Health) from the midline and the pia to the center of the lesions and aligned to 4 matching coronal slices from the Mouse Brain Atlas (Allen Institute) for AP axis reference.

### Electrophysiology and Data Acquisition

Electrophysiology signals were acquired continuously at 30 kHz, while the behavioral tracking was acquired at 30 Hz with one or two lighthouse tracking stations (HTC Vive Base Station, Amazon). An additional camera (FFY-U3-16S2M-S, FLIR) was placed on the ceiling of the behavior box for behavior monitoring. 3 mice were recorded on a first generation Open-Ephys system (Siegle, et al., 2017) with an Intan 64 or Intan 32 (for the carbon fiber-implanted mouse) headstage. In these mice, tracking provided by two lighthouse receivers (TS4231, Digikey) attached at the base of the headstage, whose signal was recorded and powered through a teensy 3.6. The other 9 mice were recorded on a new-generation Open-Ephys ONIX (Newman, et al., 2019) system with 64 channel headstages with a powered commutator, that integrated electrophysiology and behavior tracking using the Bonsai software (Lopes, et al., 2015).

Spikes were sorted on 300-6000Hz band pass filtered continuous traces, using MountainSort (https://github.com/flatironinstitute/mountainsort, Chung *et al*., 2017). Units were then manually selected based on the spike template shapes and interspike interval (ISI) distribution. After implant surgery, ADn-targeting tetrodes were lowered until HD-coding cells were identified based on their tuning obtained from brief recordings, and RSC targeting tetrodes were slowly lowered until well-isolated units appeared. When at least 3 HD cells in ADn were first detected, recordings of cue rotation or cue-on-off sessions were collected over a minimum of 2 weeks and up to 8 months. Tetrodes in ADn and RSC were regularly moved by ∼20-40 um increments, followed by recordings of short stable cue-on sessions to verify if the yield was improved. To avoid sampling of the same units for HD neurons quantifications and spike correlations for monosynaptic connections, sessions recorded at least 4 days apart and only one session for the carbon fiber-implanted mouse were included in these analyses. At the end of the experiments, electrolytic lesions were obtained by passing positive and negative current (20-25 uA) on each electrode contact for 5 s with a stimulus isolator (A365RC, WPI) while the animal was under isofluorane-induced anesthesia. After 30-60 min of recovery, the brains were extracted for histology.

### Data Analysis HD unit selection

HD was quantified as the relative orientation of two or three infrared lighthouse receivers present on the integrated headstage, after their (x,y) coordinates were linearly interpolated to align to the same 50Hz timestamps. For each session, HD tuning curves were quantified as the histogram of the spike trains over HD angles of 10-degree bins divided by the occupancy. To minimize degradation of the HD tuning over cue rotations or cue on-off trials, the first trial or a period of long cue-on stability and sufficient angle occupancy was used for each session. For HD unit selection and information metrics for other spatial correlates, data from a stable cue-on period was used. Information was calculated as bits/spike as

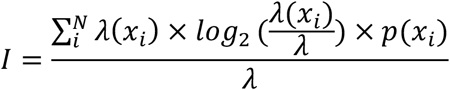

following the methods of (Skaggs, et al., 1993) where *x* is the binned HD (N=36 bins), p(x) is the occupancy, and *λ* is the mean firing rate and *λ*(*x*_*i*_) is the firing rate for each angular bin. Cells in ADn and RSC were selected as HD-coding if the amount of directional information was more than the 95^th^ percentile of the shuffle distribution and the resultant of the smoothed tuning curve was more than the 90^th^ percentile of the shuffle distribution. Shuffling of the spikes was obtained by shifting 500 times the spike trains by random amounts with respect to the HD from tracking. The peak number was obtained from MATLAB’s “*findpeaks*.*m*”, with a minimum peak distance of 1.2, width of 0.7 and prominence of 4. For units with more than one peak (Supplementary Fig. 2A) and with prominence more than 10, we applied MATLAB’s “*fitnlm*.*m*” with a basic von Mises model function with one peak

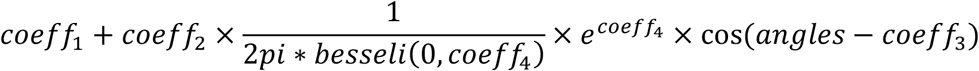

where *besseli* is the modified Bessel function of the first kind, *angles* is the range of possible angles between 0 and *2pi* in 3600 bins, and the subscripted coefficients correspond to: 1 a baseline constant offset, 2 a scaling factor for the peak height, 3 the peak location, 4 the concentration parameter in the Von Mises probability distribution. For instances where up to 3 peaks were identified, the model function was expanded with a linear sum combining additional sets of height, location and concentration coefficients. Starting values for coefficients estimation were obtained from the *findpeaks*.*m* and 0 for the constant offset. The aim of this strategy was to identify a von Mises distribution anchored to the largest peak in tuning curves, whose resultant would have otherwise been much lower despite a strong directional information (Fig. 1E 3^rd^ example from top, and Supplementary Fig. 2A). Angular velocity was calculated as the first derivative in the unwrapped HD.

### HD Decoding

We decoded HD using a linear-Gaussian GLM based on the 20 ms-binned firing rates of ADn and RSC neurons, separately (Supplementary Fig. 3A). A Butterworth filter with cutoff normalized frequency of 0.2 was applied to the firing rates, which were then normalized. Maximum a posteriori (MAP) estimation coefficients of the neuronal ensembles (the predictors) were obtained via ridge regression regularization was applied to the sine and cosine of HD, binned in 10° bins. The segments for the training were taken from a cue-on period at least 50 s away from rotation. Decoding was performed using MATLAB’s “*glmval*.*m*” with the corresponding identity link function and HD reconstructed as the 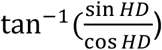from the decoded bins. With this strategy, we obtained the HD representation, which was linked to the neuronal ensembles via the learned coefficients during training, in a test period following the training session and in the period after cue rotation. For testing drift in dark and light, shorter training sessions during a stable cue-on (up to 8 min of data) were used and evaluated on the subsequent cue-on period and after the cue was turned off.

### Detection of putative functional monosynaptic connections

We performed spike cross-correlation between all unique possible pairs of simultaneously recorded ADn-RSC neurons to detect putative monosynaptic connections. Excitatory connections appear as peaks in the cross-correlogram in the short time scale (1-5 ms) above baseline (Fujisawa, et al., 2008; Stark, et al., 2009). We focused on excitatory connectivity since corticothalamic and thalamocortical projections are excitatory and negative deviations from the baseline are often result from large positive deviations in the other direction. Cross correlograms were constructed in bins of 0.5 ms by taking all spikes occurring during cue-on trials in a session (or only during cue-off trials). The baseline correlation, simulating homogeneous firing, was constructed by convolving the cross correlogram with a 10 ms s.d. Gaussian window. Significant connections were detected if at least 2 consecutive bins in the 1 to 5 ms window of the cross correlogram were above the 99.9^th^ percentile of the cumulative Poisson distribution at the baseline rate.

### Interneuron and Pyramidal neurons classification

Fast spiking (FS) interneurons and regular spiking (RS, pyramidal) neurons have distinct features that appear on the extracellular spike waveforms and can be used for classification (Barthó, et al., 2004; Wilson, and McNaughton, 1993). We applied the metrics described in (Sirota, et al., 2008) on spike waveforms identified from the bandpass filtered continuous traces. Briefly, a mean spike waveform was obtained for each cortical neuron and the peak-to-trough was quantified as the time between the peak of the spike and the maximum point in the afterhyperpolarization, whereas the symmetry around the spike was calculated as the difference between the height at the maximum point after spike peak and the maximum point before spike peak, divided by the sum of these two quantities. While in our dataset the symmetry value was unimodally distributed, the peak-to-trough was clearly bimodally distributed, allowing to cluster FS and RS with a previously reported (Peyrache, et al., 2015) cutoff duration of 0.42 ms, which resulted in average spike waveforms with a slow repolarization decay for RS and faster repolarization in FS (Supplementary Fig. 7).

### Quantification and Statistical Analysis

All statistical analyses were performed in MATLAB (MathWorks, R2020a). All spiking and behavioral data, with exception of the spike times for the detection of monosynaptic connections, was binned in 20 ms bins. Behavioral tracking from the light receivers was linearly interpolated. Circular Statistic Toolbox (Berens, 2009) functions were employed for quantifications of HD units’ preferred firing directions (PFD) as circular means, as well as neural population and decoded errors mean rotations, confidence interval, and circular correlations between regions, preferred direction difference of HD cell pairs between trials, and drifts between regions. Correlations between before and after rotations HD units tuning curves and between the absolute egocentric bearing of the cue and the size of the rotation were calculated as Pearson correlations coefficient.

Angular velocity (AV) modulation during cue-on was calculated for the same HD neurons identified in Fig. 1C as the ratio of the firing rate for high angular velocity (>30°/s) over the firing rate for low angular velocity(<30°/s) taken from the entire session away from cue rotations. Data in Fig. 1G-I and Supplementary Fig. 2F and top of H also came from the same sessions, but only trials where significant tuning curves rotations were observed and where each direction bin had a 1s minimum occupancy. For cue-off angular velocity modulation, HD neurons from sessions with dark periods recorded at least 4 days apart after tetrode advancement were included. The same sessions were used to calculate the change in HD score between cue-on and cue-off trials and only trials were all HD bins were occupied for at least 1 s were included.

The relationship between the tuning curve correlations before versus after rotations and the change in preferred direction was fitted using “*fitnlm*.*m*” with robust weighing option to estimate the parameters with a gaussian model function:

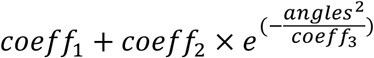

Two-tailed Kolmogorov Smirnov tests were used to compare the distribution of HD score changes in ADn and RSC and of the time lags of peak correlation between ADn and RSC decoded errors vs the null distribution obtained from 100 shuffles, and Wilcoxon Signed-Rank tests to compare the peak correlation values before and after cue rotation. Shuffle distributions for the decoded rotations in ADn and RSC (Supplementary Fig. 3F&G) were obtained by decoding the HD from circularly shifted spikes by random amounts 100 times. Shuffle distributions for the decoded errors (both for dark and cue-rotation drifts) were obtained by circularly shifting the tracked HD by random amounts 100 times and subtracting it from the RSC decoded HD. Significant ensemble preferred directions rotations were determined if at least half of the HD cells experienced preferred direction shifts larger than the 98^th^ percentile of a distribution obtained by randomly reassigning 500 times the indices around that rotation trial. Unless otherwise stated, summary data was presented as mean and 95% confidence intervals and P-value threshold of 0.05 were used for statistical non-parametric tests. Multiple comparison tests were performed with Bonferroni-correction.

## Acknowledgements

We thank Hongkui Zeng, Shenqin Yao, Ali Cetin, and the Allen Institute as well as Ian Wickersham and Heather Sullivan for sharing monosynaptic rabies tracing viral constructs. We thank Mila Halgren and Lukas Fischer for feedback on the manuscript and members of the Harnett laboratory and Wisam Reid for constructive criticism on the project. This work was supported by a MathWorks Graduate Fellowship (M.S.H.v.d.G), NIH K99 6943778 (J.V.), RO1NS106031 (M.T.H.), the James W. and Patricia T. Poitras Fund at MIT (M.T.H.), and the Klingenstein-Simons Fellowship Program (M.T.H).

## Author Contributions

M.S.H.v.d.G and M.T.H. conceived the project with input from J.V. M.S.H.v.d.G designed and performed all experiments and analyses, and wrote the manuscript with M.T.H. and J.V. J.V. and J.P.N. provided guidance with the Open-Ephys and ONIX acquisition systems. J.V. provided guidance in surgery, drive fabrication and, together with E.H.T., suggestions in data analysis. N.J.B. helped with histology and P.V. with preliminary analyses. M.T.H. supervised all aspects of the project.

