## Supplementary Figures and Legends for "Coordinated Head Direction Representations in Mouse Anterodorsal Thalamic Nucleus and Retrosplenial Cortex"

### Supplementary Information

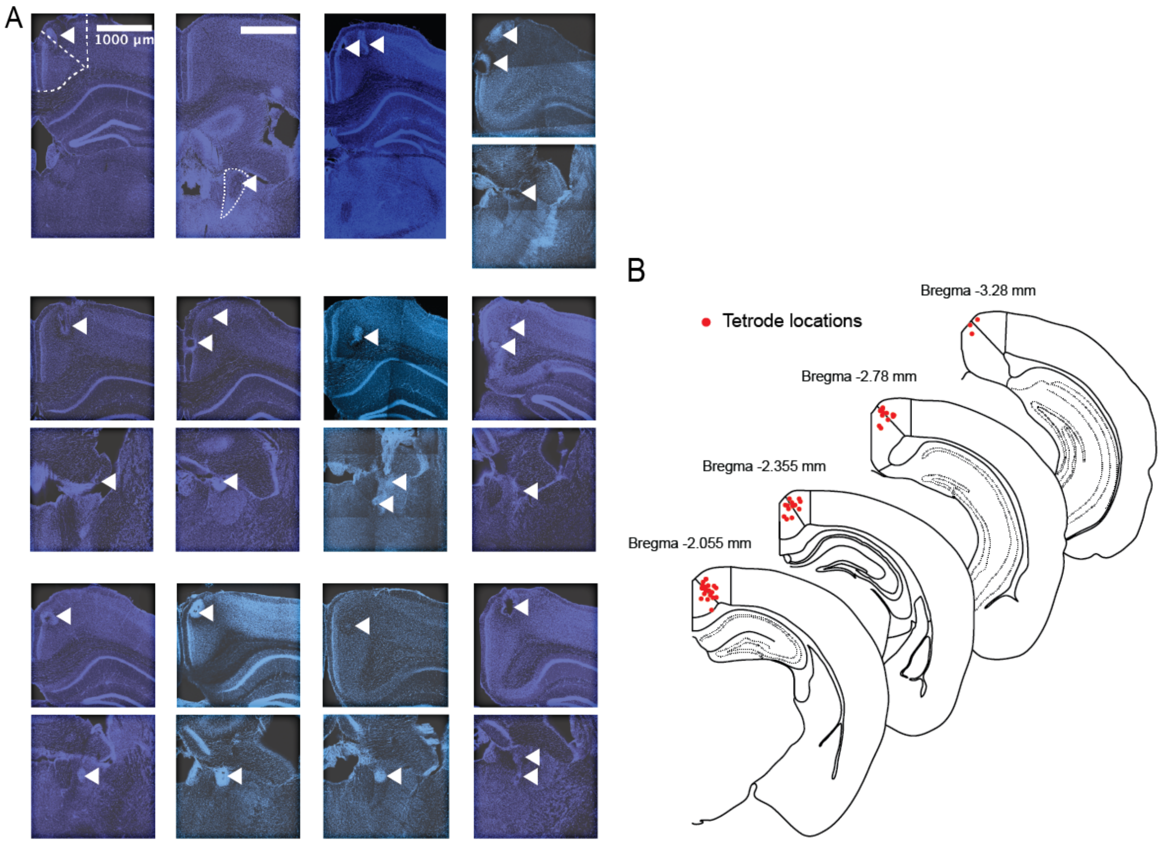


**Supplementary Fig. 1: Electrolytic lesions in RSC and ADn A:** Confocal images of coronal slices containing ADn or RSC from individual mice used for tetrode recordings. Images from individual mice are organized in a 3x4 grid, where sample electrolytic lesions in RSC are presented in the top row and those in ADn in the bottom row for mice with dual site recordings. For mice with single recording site (the first 3), only one sample image is provided (in order RSC, carbon fiber-targeted ADn and RSC). ADn and RSC granular and dysgranular outlines are shown with dashed lines in the first two images. White, leftward arrows indicate the lesions, which corresponds to the locations of the recording electrodes. Scale bar 1 mm; the same scale was applied to all images. **B:** Summary of cortical tetrode locations from all mice, distributed over 4 roughly matching coronal slices from the Mouse Allen Brain Atlas.

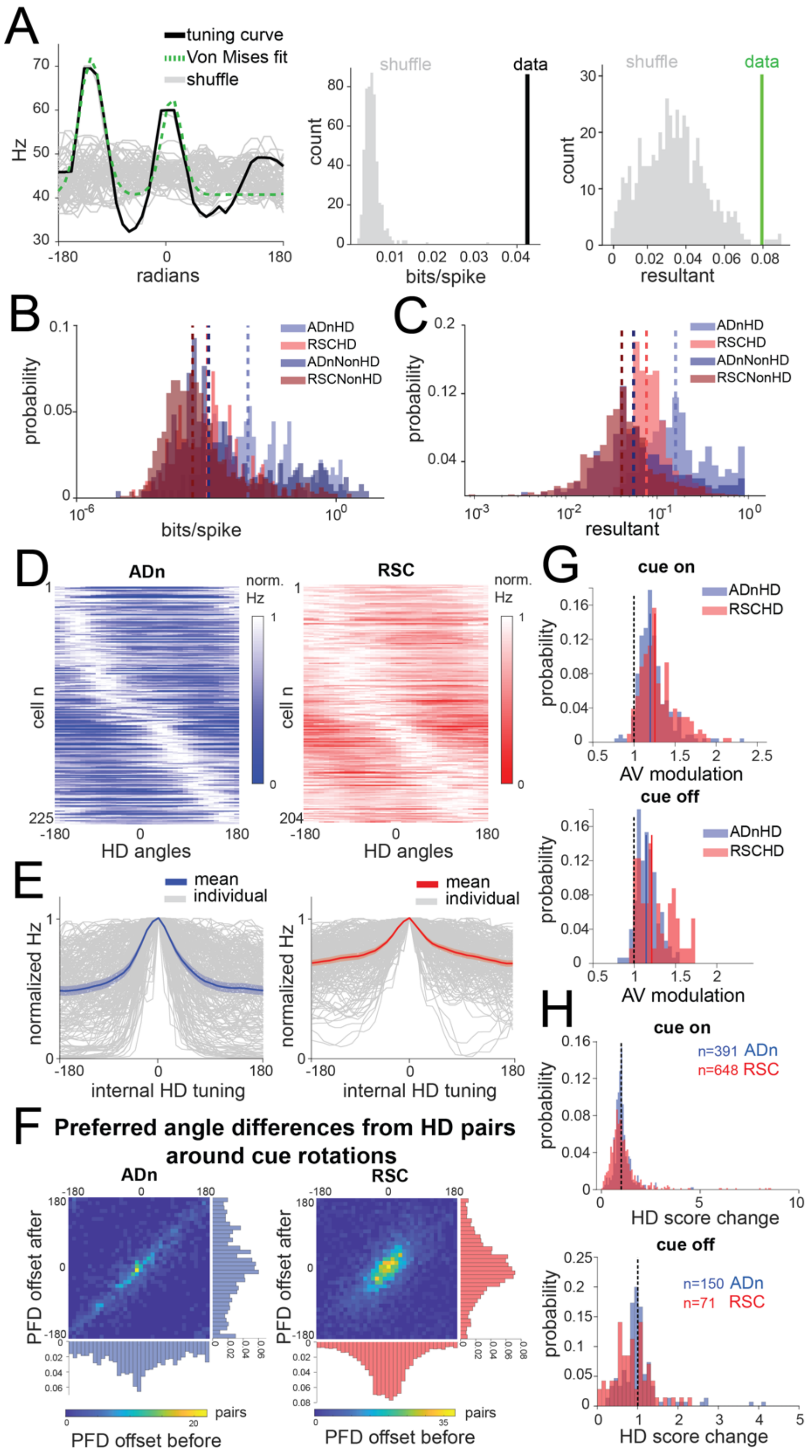


**Supplementary Fig. 2: HD selection method and HD features across ADn and RSC**
**A:** Left, example of von Mises fitting (green dashed) to the tuning curve (solid black) with two identified peaks in ADn, compared to the tuning curves obtained from shuffling the spikes (grey solid lines). Middle, the directional information calculated from the tuning curve in the example on the left is above the 95$^{th}$ percentile of the shuffle distribution. Right, the corresponding resultant of the fit to the largest peak of the tuning curve is above the 90$^{th}$ percentile of the shuffle distribution. **B:** Distributions and medians (dashed lined) of the directional information (in

bits/spike) across ADn HD (0.0567, n=225) and non-HD (0.0142, n=248) and RSC HD (0.0134, n=205) and non-HD (0.0074, n=805) units (p<0.0001, Kruskal Wallis test, and p<0.001 for multiple comparisons after Bonferroni correction, except for ADn NonHD and RSC HD cells, where p=0.0675). **C:** Distributions and medians (dashed lines) of the resultants of the same groups of units as in B: ADn HD (0.159, n=225) and non-HD (0.055, n=248) and RSC HD (0.076, n=205) and non-HD (0.041, n=805) units (p<0.0001, Kruskal Wallis test, and p<0.001 for multiple comparisons after Bonferroni correction). **D:** Normalized tuning curves of all the HD units in Fig.1c sorted by peak location in ADn (left) and RSC (right). **E:** Individual (grey) and mean and 95% CI of normalized and peak-aligned to 0° HD tuning curves from D, ADn (left, blue) and RSC (right, red). **F:** Unnormalized heatmaps of the 2D histograms shown in Fig. 1G with marginal distributions of the preferred direction (PFD) differences between HD pairs before and after rotations. **G:** Distributions and medians of angular velocity (AV) modulation of HD units firing in ADn (blue) and RSC (red) during cue-on (top, ADn n=225, RSC n=204) and cue-off conditions (bottom, ADn n=150, RSC n=57). In both conditions RSC AV modulation was more prominent than ADn (Mann-Whitney test p=0.0007 and p=0.0033 for cue-on and cue-off, respectively). **H:** Distributions of HD units score ratios of ADn (blue) and RSC (red) for cue-rotations trials (top) and cue on-off trials (bottom). For both manipulations, only a subset of sessions that were at least 4 days apart to ensure sampling of different units and only trials were each HD bin was occupied for at least 1 s were included. For the cue rotation trials, only trials with at least 2 HD units and with a significant rotation were included. Cue-rotations: ADn n=391, median=0.99, RSC n=648, median 0.94, Kolmogorov-Smirnov test p<0.0001. Cue on-off: ADn n=150, median=0.97, RSC n=71, median 0.82, Kolmogorov-Smirnov test p<0.001.

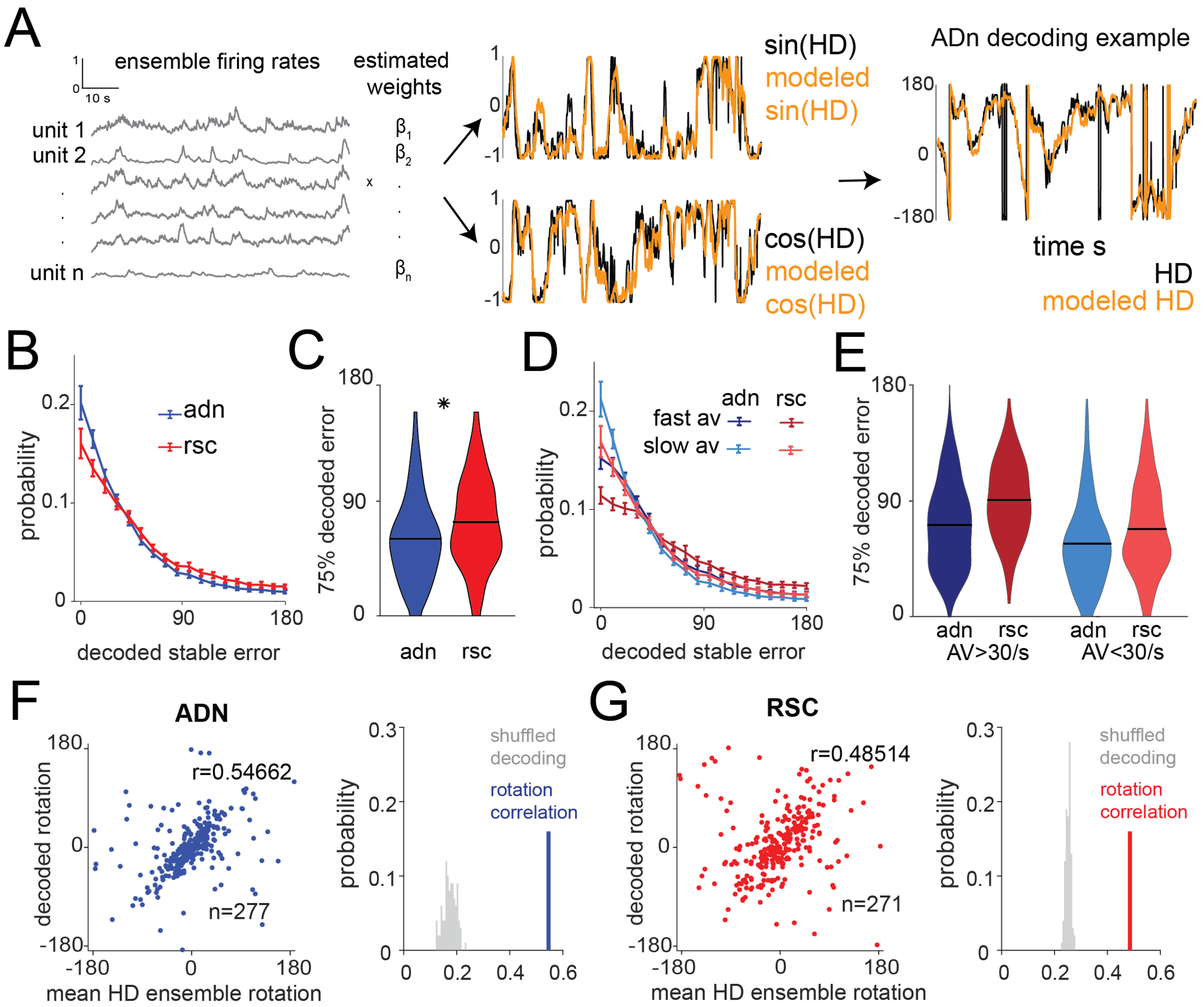


**Supplementary Fig. 3: HD decoding with a linear-Gaussian GLM. A:** Linear-Gaussian GLM-based decoding strategy for each trial and neuronal ensembles (See also Methods Section) with an example of modeled sin(HD) and cos(HD). **B:** Mean and 95% CI of the absolute decoding error distribution for ADn (blue) and RSC (red) from the test period for all trials (n=303 for ADn, n=311 for RSC). **C:** Violin plots of the $75^{th}$ percentile of the absolute decoded error distributions for ADn and RSC (p<0.0001, Mann-Whitney test, means highlighted in black 60.133° in ADn, n=303, and 72.46° in RSC, n=311). **D:** Same as B, but separating time points of high (>30°/s) from low (<30°/s) AV. **E:** Same as C, but separating between the 75% errors collected from high versus low AV (means of low AV: ADn=56.8°, RSC=68.46°, means of high AV: ADn=71.67°, RSC=90.83°; 2-way ANOVA p<0.0001 between the two regions and between the two velocity states, p=0.045 of the interaction between the two groups). P<0.001 for multiple comparisons test between all groups except high AV ADn and low AV RSC (p>0.05). **F:** Left: scatter plot of the mean rotations from simultaneous ADn HD neurons tuning curves versus the mean rotations calculated from decoding HD from ADn neurons (circular correlation coefficient=0.54, p<0.0001, n=277 trials). Right: the observed correlation is more than the $99^{th}$ of 100 correlation obtained from shuffling the spikes for HD decoding. **G:** Same as F but for RSC (circular correlation coefficient=0.48, p<0.0001, n=271 trials).

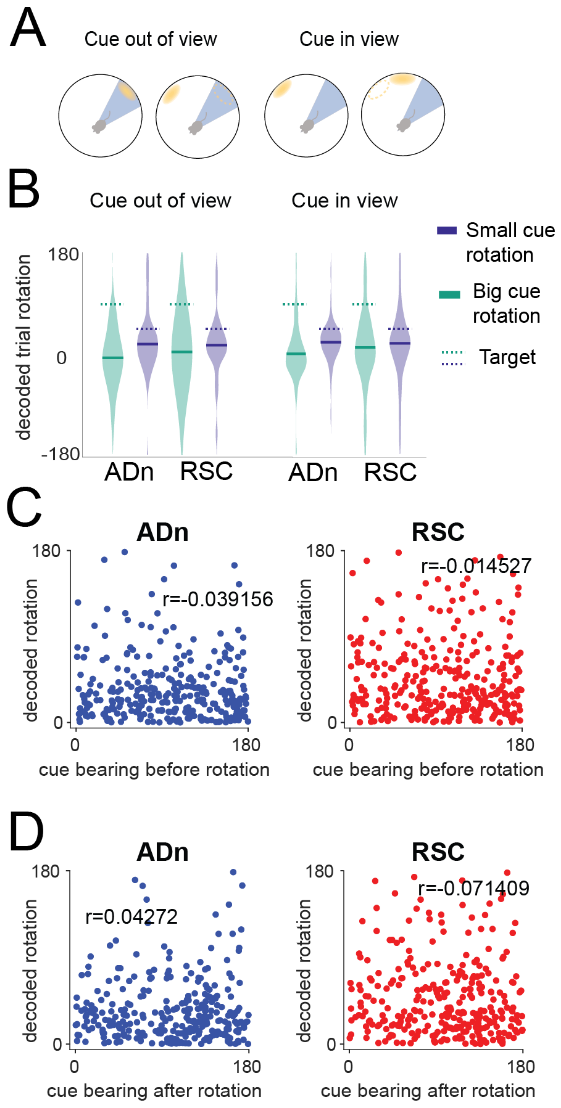


**Supplementary Fig. 4: Relationship between cue bearing and decoded neural rotations** **A:** Schematic of the distinction between egocentric "in view" (right) and "out of view" (left) angular position of the visual cue before rotation. **B:** Violin plots of the decoded rotations from simultaneous ADn and RSC trials according to the imposed cue rotation size (at a cutoff of 67.5°) and the egocentric view of the cue before rotation. Large cue rotations resulted more often in neural rotations around 0 than the small cue rotations (multiple ways ANOVA, p=0.010 effect of the size of rotation, p>0.05 for effects of brain region and egocentric condition and for all interactions. P>0.05 for all Bonferroni-corrected multiple comparisons, between the following categories N trials and means: large/out of view: n=18, ADn=-0.127, RSC=0.0569, large/in view: n=100, ADn=-0.001, RSC=0.194; small/out of view: n=26, ADn=0.31, RSC=0.26; small/in view: n=69, ADn=0.35, RSC=0.32). **C:** Decoded rotation as a function of the egocentric bearing of the cue before rotation. **D:** Decoded rotation as a function of the egocentric bearing of the cue after rotation. P>0.05 of the Pearson's correlation coefficients, ADn n=300, RSC n=306, both in C and D.

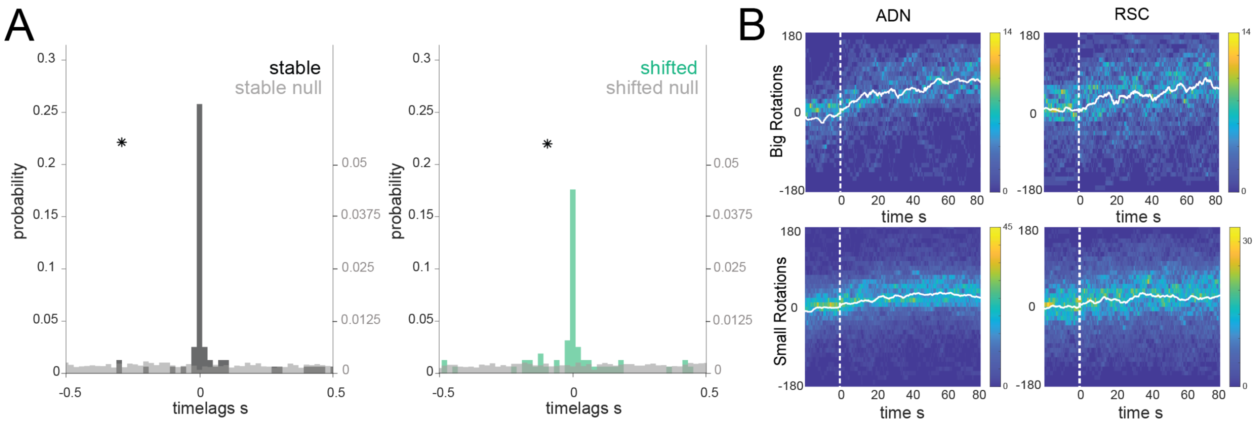


**Supplementary Fig. 5: Time course of the decoded HD rotations. A:** Similar to Fig. 2E but for only the first 25s of simultaneous decoded errors after cue rotation. As in 2E, asterisks indicate two-sample Kolmogorov-Smirnov test, $p<0.0001$ for both stable and shifted versus the null distributions in grey. No difference between stable and shifted trial correlations was observed (Wilcoxon Signed-Rank test $p=0.38$). **B:** 2D heatmaps of the decoded errors (in 10° bins, y axis, over 20 ms bins, x axis) from 20 s before the rotated cue entered the visual field of the mouse up to 80 s after. All traces were aligned to have final positive target offsets; for simultaneous RSC-ADn trials the target was based on the mean of the ADn decoded errors. Large rotations (offset larger and equal than 67.5°) ADn n=46, RSC n=72; small rotations (offset smaller than 67.5°and larger than 11.5°) ADn n=182, RSC n=185. White lines indicate the circular means across the trials.

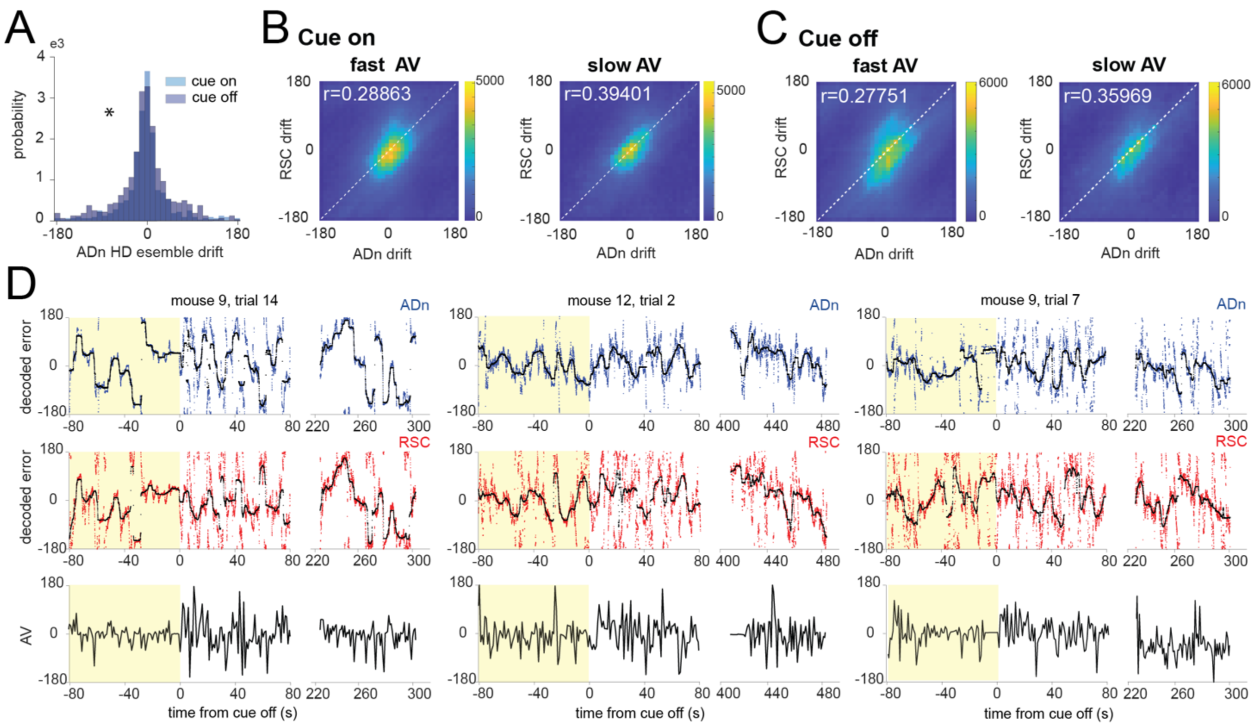


**Supplementary Fig. 6: Effect of angular velocity (AV) on HD drift in darkness in ADn and RSC. A:** Distribution of the ADn HD ensemble mean of the differences between the preferred directions from the first 4 minutes of cue-on and either the subsequent cue-on (lighter shade) and cue-off (darker shade) periods calculated every 2 minutes ($p<0.001$ Kuipers test, 72 trials, 5 mice). **B:** Heatmap of 2D histograms of 10°-binned ADn vs RSC decoded HD errors (referred to as drift) from time points of fast AV (>30°/s, left) and slow AV (< 30°/s, right) while the cue was on. Consistent with a lower decoded accuracy (Supplementary Fig. 3C) and the effect of AV on HD firing rate (Supplementary Fig. 2F), the circular correlation coefficient is lower for high AV. **C:** Same as B but for cue-off periods. **D:** Examples of decoded HD representations from different cue-on/cue-off trials of simultaneously recorded ADn (blue) and RSC (red) during cue-on (yellow shade) and cue-off periods. In black overlaying, the median-smoothed decoded errors over a 5 s window. Bottom plots, the corresponding AV profile over time.

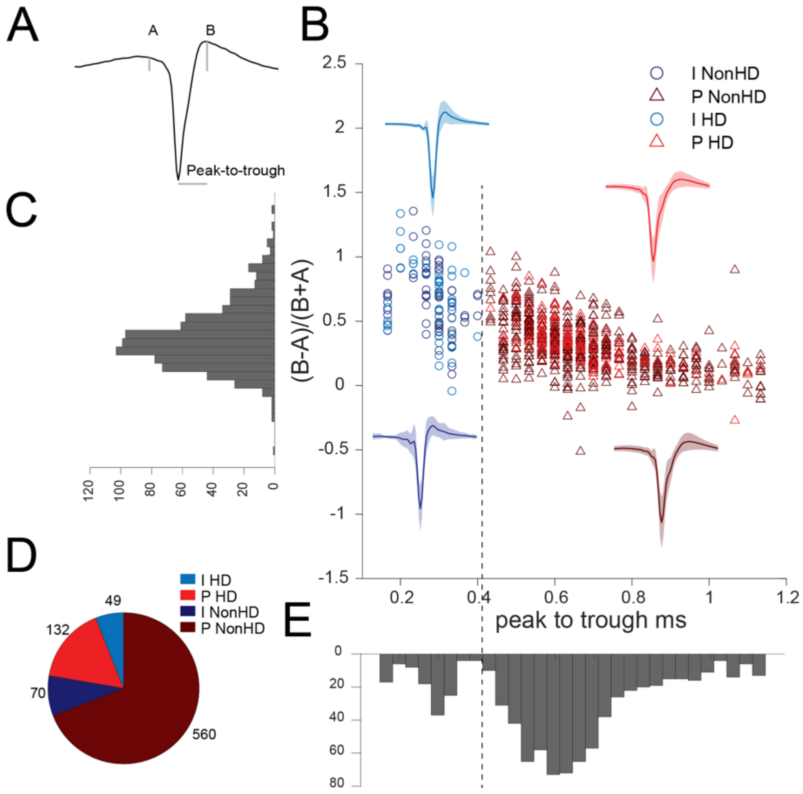


**Supplementary Fig.7: Separation of putative pyramidal neurons and fast-spiking interneurons in cortex. A:** Example putative RS neuron spike waveform with A (pre-polarization) and B (after hyperpolarization) heights and the peak to trough distance metrics used for separating RS (~pyramidal) and FS (~interneurons) spiking neurons in RSC. **B:** Scatter plot of all 811 RSC units according to the metrics calculated as in A. Dashed line indicates the <0.42 ms peak-to-trough duration discrimination. Units are shape- and color-coded to reflect the unit type classification and whether they were HD tuned or not. The plot includes the mean and SD of the combined spike waveforms for the 4 classes of units. **C:** Unimodal distribution of the spike waveform symmetry values across all units. **D:** Counts of the 4 classes of units. E: Bimodal distribution of the peak-to-trough values, indicative of two clusters.
